# Probing SWATH-MS as a tool for proteome level quantification in a non-model fish

**DOI:** 10.1101/2020.02.23.946913

**Authors:** Alison A. Monroe, Huoming Zhang, Celia Schunter, Timothy Ravasi

## Abstract

Quantitative proteomics via mass spectrometry can provide valuable insight into molecular and phenotypic characteristics of a living system. Recent mass spectrometry developments include data-independent acquisition (SWATH/DIA-MS), an accurate, sensitive, and reproducible method for analyzing the whole proteome. The main requirement for this method is the creation of a comprehensive spectral library. New technologies have emerged producing larger and more accurate species-specific libraries leading to a progressive collection of proteome references for multiple molecular model species. Here, for the first time, we set out to compare different spectral library constructions using multiple tissues from a coral reef fish to demonstrate its value and feasibility for non-model organisms. We created a large spectral library composed of 12,553 protein groups from liver and brain tissues. Via identification of differentially expressed proteins (DEPs) under fish exposure to environmental stressors we validated the application and usefulness of these different spectral libraries. Successful identification of significant DEPs from different environmental exposures occurred using the library with a combination of DIA+DDA data as well as both tissue types. Further analysis revealed expected patterns of significantly upregulated heat shock proteins in a dual condition of ocean warming and acidification indicating the biological accuracy and relevance of the method. This study provides the first reference spectral library for a coral reef fish and for a non-model organism. It represents a useful guide for the future building of accurate spectral library references in non-model organisms allowing the discovery of ecologically relevant changes in the proteome.

## Introduction

Proteomics provides insight into complex biological mechanisms and cellular phenotypes by measuring the presence and abundance of proteins controlling the execution of molecular processes (Aebersold & Mann, 2016; Y. Liu, Beyer, & Aebersold, 2016). The possible post-translational modifications inferred from proteomics can have a stronger correlation to phenotypic observations than transcriptomics, although the technology to quantify the proteome lags behind that of the transcriptome (Y. Liu et al., 2016; Tang et al., 2015). In the past decade proteomic techniques have been developing rapidly, especially those dealing with quantitative proteomics. Previously, quantitative methods were defined in two categories: shotgun and targeted, each with different strengths and weaknesses. The shotgun method identifies more peptides, but has reduced quantitative accuracy and reduced reproducibility due in part to the necessity for prefractionation prior to liquid chromatography mass spectrometry (LC-MS) analysis (Michalski, Cox, & Mann, 2011). These methods also require costly chemical labelling that increases data processing times. Targeted proteomics (S/MRM), on the other hand, is better for reproducibility if the proteins in question are known. However, this method is limited in the number of measurements and therefore peptides found (Gillet et al., 2012).

Sequential window acquisition of all theoretical spectra (SWATH/DIA-MS) is a newer method that combines the strengths of shotgun and targeted proteomics (Gillet et al., 2012). SWATH/DIA-MS is a label free and relatively cheap mass spectrometry method using data independent acquisition (DIA) combined with proteome wide spectral libraries created from data-dependent acquisition (DDA) methods against which the fragment ion maps of the DIA method are aligned (Gao et al., 2017; Gillet et al., 2012; Huang et al., 2015). Using the sequential isolation window acquisition has achieved high fragment ion specificity and the ability to identify and quantify thousands of proteins in one measurement (Gillet et al., 2012; Tang et al., 2015). Recent studies have also shown high reproducibility of results when using the same cell cultures across different labs as well as different computational software (Collins et al., 2017; Navarro et al., 2016). Most SWATH/DIA-MS studies to date have used model organisms such as mice, humans, zebrafish, or yeast with high quality genomic resources and little biological variation to great success (Blattmann et al., 2019; Braccia, Espinal, Pini, De Pietri Tonelli, & Armirotti, 2018; Bruderer et al., 2015; Collins et al., 2017; Krasny et al., 2018; Rosenberger et al., 2014; Zhang et al., 2019). A recent study has looked into the heat stress response of broiler chickens (*Gallus gallus domesticus*) and found significant changes across the proteome in ecologically relevant proteins with ties to heat stress responses (Tang et al., 2015). Despite this being a non-model organism, over generations it was kept in controlled conditions and lacks the natural variation of wild organisms. The above qualities of SWATH-MS suggest that it is an ideal method to quantify peptides at the proteome level across large numbers of biological samples with high quality reproducible results, and also an impactful way to understand complex mechanistic and phenotypic differences in particular in non-model organisms which usually lack genomic and proteomic resources. Currently, however, it is still unknown how effective it will be at identifying protein expression differences in wild populations with intrinsic individual variation.

The key to the SWATH/DIA-MS method is the creation of a high-quality spectral library against which to measure peptide quantities in the DIA data of the samples of interest. This reference can provide a standardized set of protein identifications making it easier to compare proteomes across experiments and laboratories (Blattmann et al., 2019; Rosenberger et al., 2017). Although study-specific libraries are easier to create, in the long run a large spectral library is more beneficial to the future studies of an organism by reducing the cost of sample preparation and measurement time. However, it has been understudied whether any loss of information comes from using a study-specific versus a general spectral library. Larger libraries require more stringent cut-offs to control the false discovery rate (FDR) that could lead to a loss of ecologically relevant information, while study-specific libraries could restrict identification at the proteome level due to the smaller reference (Blattmann et al., 2019; Rosenberger et al., 2017).

The SWATH/DIA-MS method provides the technological advances needed to study the whole proteome of non-model organisms. Due to the high biological variability of individuals from non-model or even wild organisms a greater number of biological replicates are required to establish statistical significance when studying changes in expression (Todd, Black, & Gemmell, 2016). This method’s reductions in cost and sample preparation time allow for the quantification of that required high number of individual samples. Also, the creation of a reference library erases the need for all compared samples to be injected in the mass spectrometer as one, creating the ability to compare a larger number of individuals and/or treatments. Due to this increased sample size and the proteome-wide exploration we may be able to determine ecologically relevant differences in protein expression both at the individual and population levels for any organism. In order to examine the usefulness of different SWATH libraries we turned to a commonly used non-model fish *Acanthachromis polyacanthus*. Most research in the past decade has focused on its behavioral and physiological changes under different environmental conditions, with a more recent focus on transcriptomic and epigenetic modifications (Jarrold & Munday, 2018; Schunter et al., 2016; Welch, Watson, Welsh, McCormick, & Munday, 2014). To date, the majority of the molecular studies on coral reef fish have focused on the liver and brain tissue due to the physiological effects on metabolism and aerobic scope under heat stress, and the behavioral effects under ocean acidification respectively (Bernal et al., 2018; Schunter et al., 2016; Veilleux et al., 2015). Only one previous study has measured the changes in protein expression: the method used was iTRAQ labelled shotgun proteomics of pooled samples hence not allowing for the comparison between many individuals or treatments (Schunter et al., 2016). The creation of a SWATH library for this fish will increase ecologically relevant proteomic studies by cutting the cost, reducing overall preparation and quantification times, and allowing for accurate and reproducible measurements of the proteome across high numbers of individuals.

Here we created several spectral libraries to evaluate the performance of a study-specific versus a large species-specific library for the non-model fish *A. polyacanthus*. We focused on liver and brain tissues due to previous studies demonstrating the effects on these tissues by climate change stressors. Recent approaches have created spectral libraries composed of the combined DDA runs, previously used to create the libraries, with the experiment-specific targeted DIA runs (Gandhi et al., 2017). Our aim is to investigate the utility of this new combined library method in the identification of peptides via targeted DIA-MS, using a complex experimental design of fish exposed to multiple climate change stressors. This manuscript provides a first reference library in a non-model fish species, as well as a guide on the efficiency, cost-effectiveness, and utility of this method in creating future proteomics references in non-model organisms aiming to evaluate genome-wide and ecologically relevant differential protein expression.

## Methods

### Fish Rearing and Tissue Dissection

*Acanthochromis polyacanthus* offspring were reared in several different conditions (Supplementary Figure 1). These include control conditions (29°C, 400μatm), elevated pCO_2_ (750μatm or 1000μatm), elevated temperature (31°C), and combined elevated temperature and elevated pCO_2_ (31°C, 1000μatm) (Jarrold & Munday, 2018; Welch et al., 2014). All experiments were undertaken at James Cook University following the university’s animal ethics guidelines (Ethics committee permits: A1828, A2210). Fish were euthanized between 3-5 months of age. Brain and liver tissues of the fish were dissected out and snap frozen in liquid nitrogen, then kept at −80°C for further processing.

### Protein extraction and digestion

Total protein was extracted using the Qiagen All prep mini kit (Qiagen) after elution through the RNA spin column. The flow-through liquid was transferred to a new Eppendorf tube with 3.5 μl of Halt protease inhibitor cocktail (Thermo Fisher Scientific) and kept on ice. The liquid was then split in half between two Eppendorf tubes (∼250 μl in each) and 1000 μl of cold acetone was added to each tube. After 30 minutes on ice all tubes were centrifuged at 4°C for 10 minutes. The samples were then moved to the fume hood and all the liquid was removed via pipetting. Samples were left to dry for 10 minutes and the resulting protein pellet was stored at −80°C for further processing. Protein pellets were resuspended in 8 M urea buffer combined with protease inhibitor (Promega) and purified using the chloroform-methanol precipitation method (Wessel & Flügge, 1984). The resulting pellet was then resuspended in 8 M UA buffer (8 M urea in 0.1 M Tris/HCl pH 8.5) and sonicated. From this, the protein quantity was measured using the Micro BCA protein assay kit (Thermo Fisher Scientific) and a SpectraMax microplate reader.

For the spectral library preparation 800 μg of total protein were combined from across all experimental conditions. This was done separately for liver and brain samples. During individual sample quantification 20 μg of protein extract from each biological replicate was used for both liver (n = 47) and brain (n = 49) samples. Protein alkylation, reduction, and digestion were done using the Filter Aided Sample Preparation (FASP) protocol (Wisniewsk et al., 2009). Following digestion with trypsin, samples were desalted using C18 filter pipette tips (Agilent) or a reversed-phase C18 Sep-Pak cartridge (Cat. WAT023590; Waters Corp.) containing an oligo R3 reversed-phase resin (Cat. 1133903; Applied Biosystems) depending on protein quantity. Elution from both the C18 cartridge and pipette tip was done using 75% acetonitrile (ACN) in 0.1% trifluoracetic acid (TFA) and all samples were subsequently dried in a SpeedVac (Thermo Fisher Scientific). For individual sample quantification with DIA-MS all samples were resuspended in 15 μl of the buffer 3% acetonitrile (ACN) in 0.1% formic acid (FA). Samples were then quantified using a NanoDrop (Thermo Fisher Scientific) and the amount of buffer was adjusted to normalize the concentration amount of each sample for injection. Indexed retention time (iRT) standards (Biognosys) were added to each prepared sample prior to the mass spec run at a 3:10 ratio (v/w).

### High pH reversed phase HPLC fractionation for spectral library preparation

For the spectral library preparation, samples were resuspended in 15 μl of Buffer A (0.1% FA). All protein derived from fish samples were combined in one tube and topped up with buffer A for a total volume of 85 μl. For this and all following steps brain library and liver library preparation were done separately but using the same methods. High pH reversed fractionation was achieved by attaching a XBridge Peptide BEH C18 column (Cat. 186003570; Waters Corp.) to an Accela liquid chromatography system (Thermo Scientific) using the HPLC application. A 135-min gradient at constant 300 nL/min was designed as follows: The gradient was established using mobile phase A (0.1% FA in H_2_O) and mobile phase B (0.1% FA, 95% ACN in H_2_O): 2.1%-5.3% B for 5 min, 5.3%-10.5% for 15 min, 10.5%-21.1% for 65 min, 21.1%-31.6% B for 13 min, 31.6%-94.7% B for 6 min, 94.7% for 6 min, and 4.7% B for 15-min column conditioning. A total of 110 fractions were collected then reduced to ∼50μl using a SpeedVac system (Thermo Fisher Scientific). Fractions were pooled into 25 groups by combining different parts of the gradient and dried with a SpeedVac system (Thermo Fisher Scientific). The dried peptides were resuspended in 0.1% FA and 3% ACN in water and protein quantity measured at A_280_ via NanoDrop (Thermo Fisher Scientific). Concentrations were normalized across all fractions and iRT (Biognosys) standards were added to each fraction at a 1:10 (v/w) ratio in preparation for the mass spectrometer.

### LC/MS-MS acquisition

An Orbitrap Fusion Lumos mass spectrometer (Thermo Fisher Scientific) was attached to an Ultimate 3000 UHPLC (Thermo Fisher Scientific) for both data dependent acquisition (DDA) and data independent acquisition (DIA) analysis. Peptides were injected and eluted through a 50 cm EASY-Spray column PepMap RSLC C18 (Cat. ES803; Thermo Fisher Scientific) with a 135-min gradient at constant 300 nL/min kept at 40°C. The gradient was established using mobile phase A (0.1% FA in H_2_O) and mobile phase B (0.1% FA, 95% ACN in H_2_O): 2.1%-5.3% B for 5 min, 5.3%-10.5% for 15 min, 10.5%-21.1% for 70 min, 21.1%-31.6% B for 18 min, ramping from 31.6% to 94.7% B in 2 min, maintaining at 94.7% for 5 min, and 4.7% B for 15-min column conditioning. For introduction into the MS an electrospray potential of 1.9 kV was used and the ion tube temperature was set to 270°C. General MS settings included default charge state of 3, application mode set to standard for peptide, and an EASY-IC was used for internal mass calibration in both MS1 and MS2 ions.

For DDA analysis the MS was set to profile mode with a resolution of 120,000 (at 200 m/z) and a full MS scan (375-1400 m/z range) was obtained. Other settings included 30% RF lens, 3 seconds between master scans, ion accumulation time of 100 milliseconds with a target value of 4e5, activation of MIPS (monoisotopic peak determination of peptide), and an isolation window of 1.6 m/z for ions. The ions carrying charges from 2^+^ to 5^+^ and measured above an intensity threshold of 5 e4 and were selected for fragmentation using higher energy collision dissociation (HCD) at 30% energy. Dynamic exclusion was used after 1 event for 10 s with a mass tolerance of 10 ppm. In DIA-MS analysis a high energy collision dissociation (HCD) fragmentation method was used with a quadruple isolation window of 25 m/z. Precursor mass range was set to 400-1200 m/z for each injection. HCD was set to 30% and the mass defect was 0.9995. MS2 was run with a scan range of 350-1500 m/z, at a resolution of 30,000, maximum injection time of 100 milliseconds, and a target value of 1e6.

### Spectral library generation

DDA raw files were processed and the library generated with Spectronaut Pulsar X (Version 12, Biognosys) using default settings. A false discovery rate of 0.01 was set at all levels (peptide, protein, and peptide spectrum match (PSM)). Peptides with a minimum length of 7 and maximum of 52 were allowed, along with a maximum of two missed cleavages. Digest type was set to specific and defined as Trypsin/P. Peptide identity was achieved via sequence alignment against the *A. polyacanthus* proteome with 36,741 entries. Default library filters in Spectronaut include ion mass to charge from 300-1800 Da, as well as minimum relative intensity of 5%, and a minimum amino acid ion length of two. Three to six fragment ions per precursor peptide were used in the library and iRT calibration required an R-squared minimum of 0.8.

A second library was generated for both liver and brain by processing all DDA and DIA runs together in Spectronaut Pulsar X. All settings were the same as above including the protein database used to search for peptide identities. To create a ‘complete’ library (combining both tissues) for *A. polyacanthus* brain and liver libraries were merged using the combine library function in Spectronaut Pulsar X.

### Quantitative Analysis of biological samples

DIA data were then loaded into the Spectronaut Pulsar X (Version 12, Biognysos) software and mapped against the generated spectral libraries leading to protein and peptide identification and quantification. Default settings of the program were applied for analysis including: estimation of FDR using a 0.01 q-value cutoff for precursors and proteins; peptide quantity measurement determination using the mean of one-three best peptides; quantitation calculated using the area of extracted ion chromatogram (XIC) at MS2 level; excluding duplicate assays. Upon completion peptide quantities for all samples (n=47 liver, n=49 brain) were exported as a data matrix containing all non-redundant protein identifications for further analysis.

Data matrices were loaded into the program Perseus v1.6.2.1 (Tyanova et al., 2016). Data was filtered by using a cutoff of at least 80% valid values in at least one condition. Protein group quantities were then log2 transformed to establish a normalized distribution and missing values were inferred using the normal distribution setting in Perseus. Differential abundance between conditions was inferred statistically using a multiple-sample test (ANOVA) with a permutation based FDR at 0.05 (250 randomizations, no group preservations). Significant differences between each of the four conditions was determined using a Tukey’s post-hoc test (FDR<0.05) in Perseus. Protein names and functions were identified with the *A. polyacanthus* genome annotation.

## Results

### Spectral Library

The key to the SWATH/DIA-MS analysis is the availability of a high-quality spectral library for the organism in question. To achieve this, we used samples from an experiment covering exposure to elevated temperature and elevated pCO_2_ as well as different tissues dissected from our organism *Acanthochromis polyacanthus*. Equal amounts of protein from each condition and each tissue were used to prevent bias in the library representation. We combined a range of protein preparation protocols to ensure both better quality and quantity peptides prior to mass spectrometry. This included purification via methanol/chloroform to remove non-proteins, an FASP protocol which improved quality and yield, and desalting using cartridges with both C18 filters and R3 material to minimize protein loss. We then used the newer Orbitrap Fusion mass spectrometry platform that has been shown to identify more peptides than other systems (Zhang et al., 2019). Lastly, the new genome developed recently for *A. polyacanthus* gave us a high-quality database of protein coding genes to search against for spectral library generation.

Based on the above optimal workflow we created six different spectral libraries to test our hypothesis. In each library we were able to identify a large number of peptides and proteins, and as expected the highest identification was discovered in the combined tissue library (Figure 1). The resulting spectral libraries contained anywhere from 49,136 to 103,022 proteotypic peptides and 8,273 to 12,553 protein groups which equates to a 22% to 35% coverage of all the protein coding genes (Table 1). Consistent with full tryptic peptide properties, most identified peptides were 8-18 amino acids in length in all libraries.

**Figure 1.**
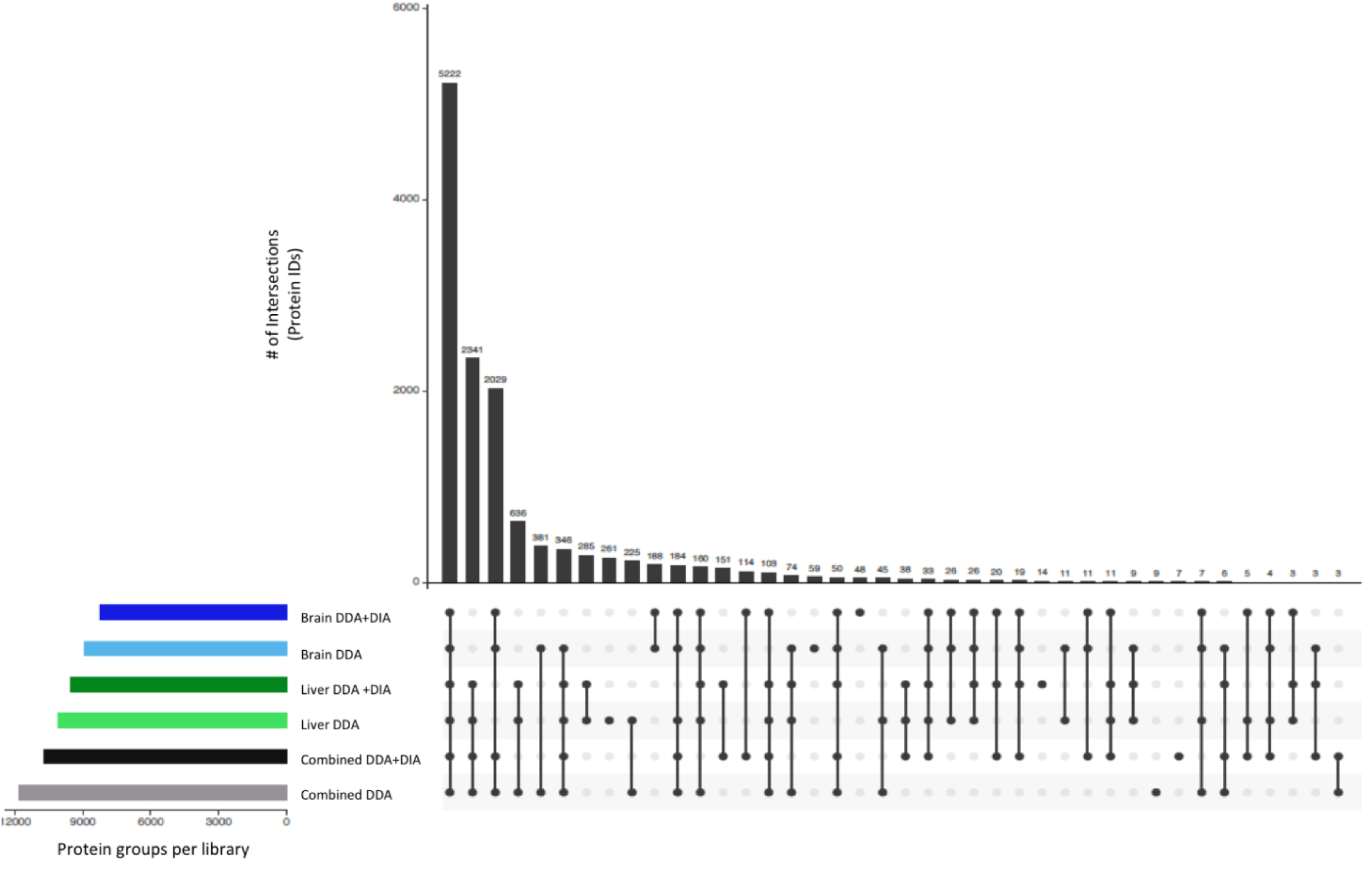
Graph of overlapping protein groups between different spectral libraries. The majority of protein identifications are all included in the combined libraries.

**Table 1.**
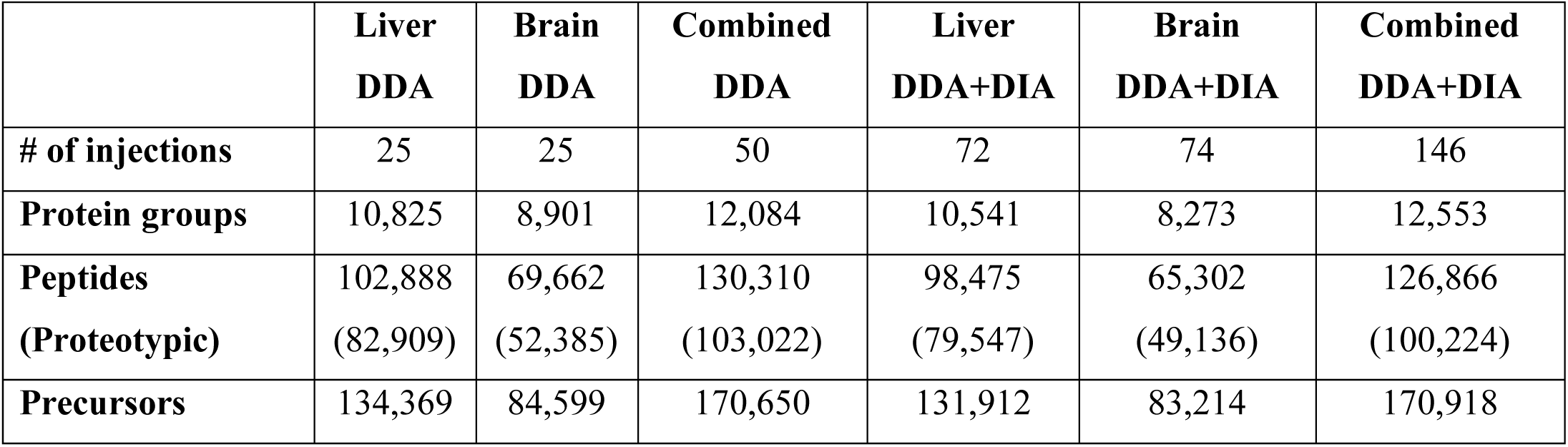
Size of the different libraries created. Proteotypic peptides are those that uniquely identify a single protein. DDA= Data dependent acquisition, DIA= Data independent acquisition

|  | **Liver**<br>**DDA** | **Brain**<br>**DDA** | **Combined**<br>**DDA** | **Liver**<br>**DDA+DIA** | **Brain**<br>**DDA+DIA** | **Combined**<br>**DDA+DIA** |
| --- | --- | --- | --- | --- | --- | --- |
| **# of injections** | 25 | 25 | 50 | 72 | 74 | 146 |
| **Protein groups** | 10,825 | 8,901 | 12,084 | 10,541 | 8,273 | 12,553 |
| **Peptides**<br>**(Proteotypic)** | 102,888<br>(82,909) | 69,662<br>(52,385) | 130,310<br>(103,022) | 98,475<br>(79,547) | 65,302<br>(49,136) | 126,866<br>(100,224) |
| **Precursors** | 134,369 | 84,599 | 170,650 | 131,912 | 83,214 | 170,918 |

### DIA/SWATH-MS for the discovery of ecologically relevant molecular processes

To determine the utility of these various spectral libraries we compared the proteomes of liver and brain tissue from *A. polyacanthus* exposed to different environmental conditions against each of the different libraries (Supplementary Figure 2). Using DDA+DIA libraries resulted in a significant increase in identified precursors and protein groups for both liver and brain samples (Figure 2). Approximately 3,300-4,100 protein groups were identified in brain depending on the library specificity and ∼2,900 to 3,100 protein groups identified in liver. A higher data completeness was revealed when using the DDA+DIA libraries versus the DDA libraries correlating to the overall number of protein groups found when using the different libraries (Table 2).

**Table 2.** Data completeness and median CV’s of both liver and brain DIA runs compared against all library types. DIA= data independent acquisition, DDA= data dependent acquisition, CV= coefficients of variance

| **Liver DIA samples** | **Data completeness** | **Median CVs** |
| --- | --- | --- |
| Liver DDA library | 50.60% | 23.1% |
| Liver DDA+DIA library | 62.60% | 31.4% |
| Brain DDA library | 54.50% | 22.2% |
| Brain DDA+DIA library | 55.30% | 25.6% |
| Combined DDA Library | 45.0% | 22.4% |
| Combined DDA+DIA library | 60.40% | 31.4% |
| **Brain DIA samples** | **Data completeness** | **Median CVs** |
| Brain DDA library | 57.30% | 15.2% |
| Brain DDA+DIA library | 77.10% | 21.2% |
| Liver DDA library | 56.7% | 14.8% |
| Liver DDA+DIA library | 56.40% | 16.8% |
| Combined DDA Library | 50.10% | 14.9% |
| Combined DDA+DIA library | 67.70% | 20.9% |

**Figure 2.**
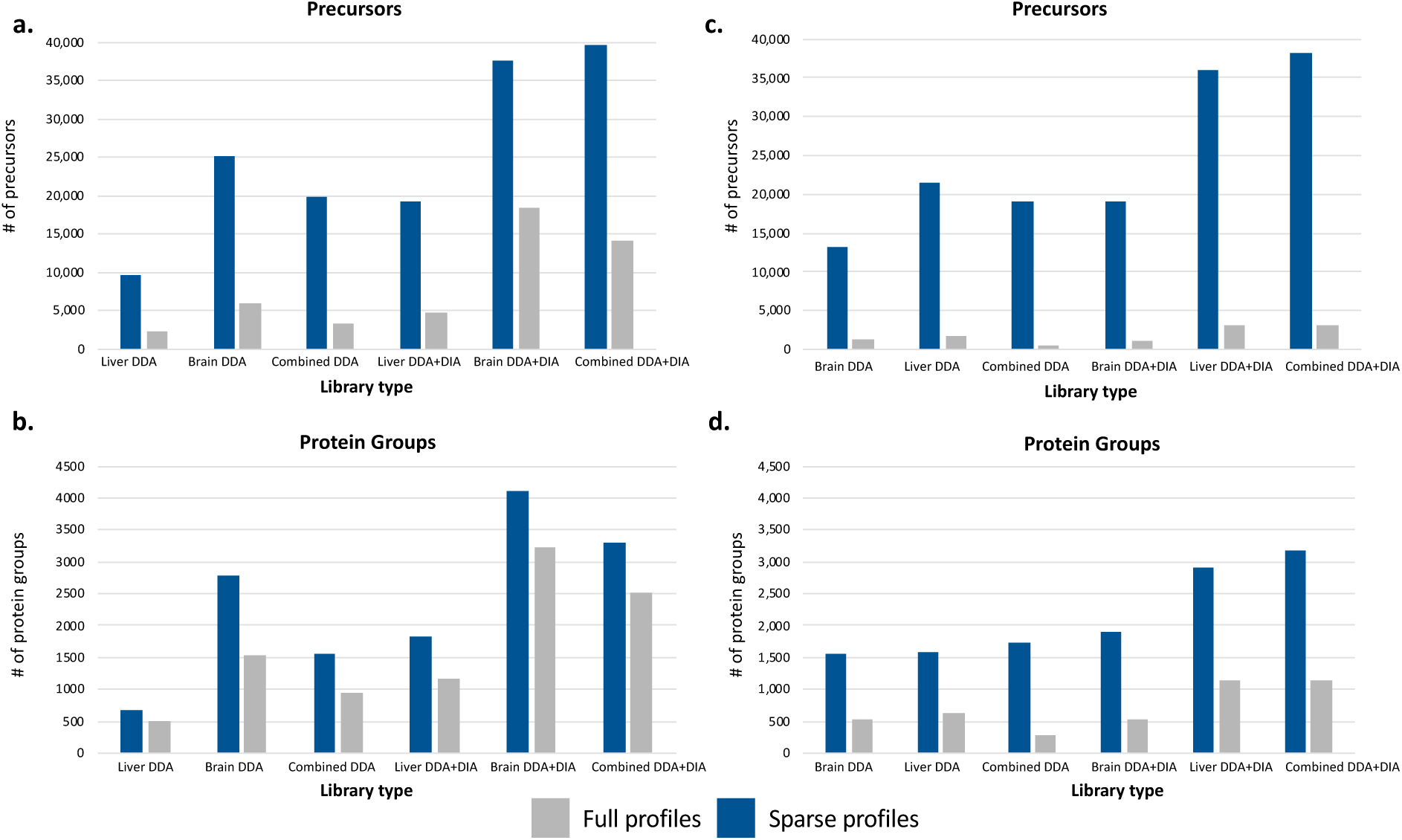
a and b represent the comparison of brain DIA samples against different libraries. c and d show comparisons of liver DIA samples from different conditions against different libraries. Full profiles indicated in gray represent proteins found across all DIA samples (liver samples are highly variable), sparse proteins represented in blue are those proteins found in at least one DIA sample.

There was little difference between the number of precursors and protein groups identified between the tissue-specific DDA+DIA library and the combined DDA+DIA library in both liver and brain. Furthering the case that a comprehensive species library combined with experiment-specific DIA runs can lead to no significant loss of identification as compared to the tissue-specific spectral library as long as it contains the targeted DIA tissue. By incorporating the DIA samples into the spectral library we are able to bolster the number of peptides represented in the library that could be found in the sample content which leads to fewer chances of false discovery without limiting the proteome coverage. To measure variability across samples and between replicates we used the median coefficient of variation (CV) that represents the ratio of the standard deviation to the mean. CVs were higher in the DDA+DIA libraries than the DDA libraries for both liver (∼31% vs 23%) and brain (∼20% vs 15%) (Table 2). Liver samples also had a small number of full profiles which refer to protein groups and precursors that are found across all DIA samples queried (Figure 2). Lack of full profiles compared to sparse profiles reveals a high variability between samples and is expected due to the high complexity of our experiment’s design.

Tissue specificity might be an issue for the proteome analysis of non-model organisms. Notably, when running the targeted DIA samples of one tissue against the spectral library of a different tissue, precursor identification decreased by an average of 43% in the liver DIA samples and ∼55% in the brain DIA samples (Figure 2). This indicates that a species-specific spectral library must contain protein samples from all targeted tissues or risk decreasing protein group identification by up to ∼75% depending on the quality of the spectral library.

### Differential Expression and functional analysis

An important function of the spectral library is to examine how well it can identify biologically and ecologically relevant differentially expressed proteins in a DIA-MS targeted analysis. Here we are able to show the definite usefulness of an experiment-specific DIA+DDA library. No unique differentially expressed proteins (DEPs) were discovered using the combined DDA library in any comparison or tissue (Figure 3). Despite having a significant amount of shared identified proteins, using the tissue-specific DDA+DIA libraries and the combined DDA+DIA libraries both identified high amounts of unique proteins (27% and 13% respectively for the brain, and 28% and 16% for the liver). This analysis made it clear that using a combined DDA library is insufficient for the identification of DEPs; when analyzing highly variable samples the library must have some specificity (tissue or study related).

**Fig 3.**
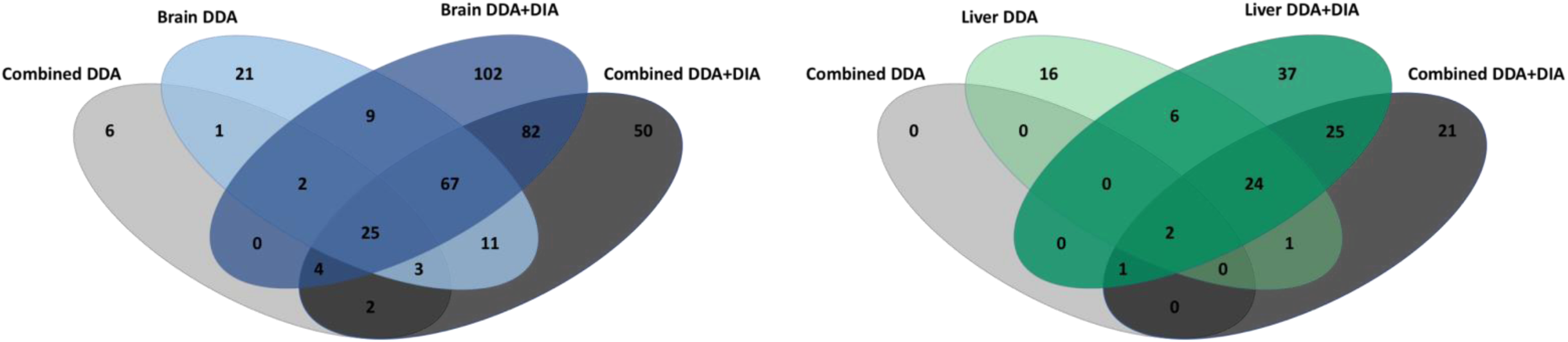
Venn diagrams of overlapping and unique differentially expressed proteins (DEPs) identified using different spectral libraries for both brain (a) and liver (b) targeted DIA analysis. Significant differential expression analyzed across all four conditions was calculated via ANOVA (FDR<0.05).

Higher numbers of statistically significant differentially expressed proteins were uniquely identified in the tissue specific DDA+DIA library in both liver and brain analysis (∼27.5% of all identified DEPs). One main goal of this study was to determine if a “whole organism” library could identify sufficient ecologically relevant DEPs as compared to a tissue-specific library. In order to determine if the reduced number of unique proteins led to any loss in ecologically relevant protein identifications, we looked deeper at the function of the DEPs identified using all library types (Supplementary Data 1). As our DIA experiment was designed to examine the stress response of a fish to dual climate change stressors (ocean warming and acidification) we examined the significant DEPs for those related to cellular stress in fish. These included proteins related to the heat shock response (HSP), a well-documented expression change seen in fish exposed to elevated temperatures (Basu et al., 2002), cytochrome related genes involved in oxidative stress, and several other previously identified stress response genes in marine fish representing protein degradation and cell death. In Figure 4 we see the combined DDA libraries provided the least identifications of the complex cellular stress response identified by the other libraries. Furthering the necessity of the inclusion of experiment-specific DIA runs in the spectral library, especially when combining tissues. In DDA+DIA libraries we see little difference between the differential expression in the combined and tissue-specific identifications (Figure 4). This suggests that using a whole organism library (including several tissues or body parts) is sufficient to identify important changes in the entire proteome when a non-model organism is exposed to varying environmental conditions.

**Fig 4.**
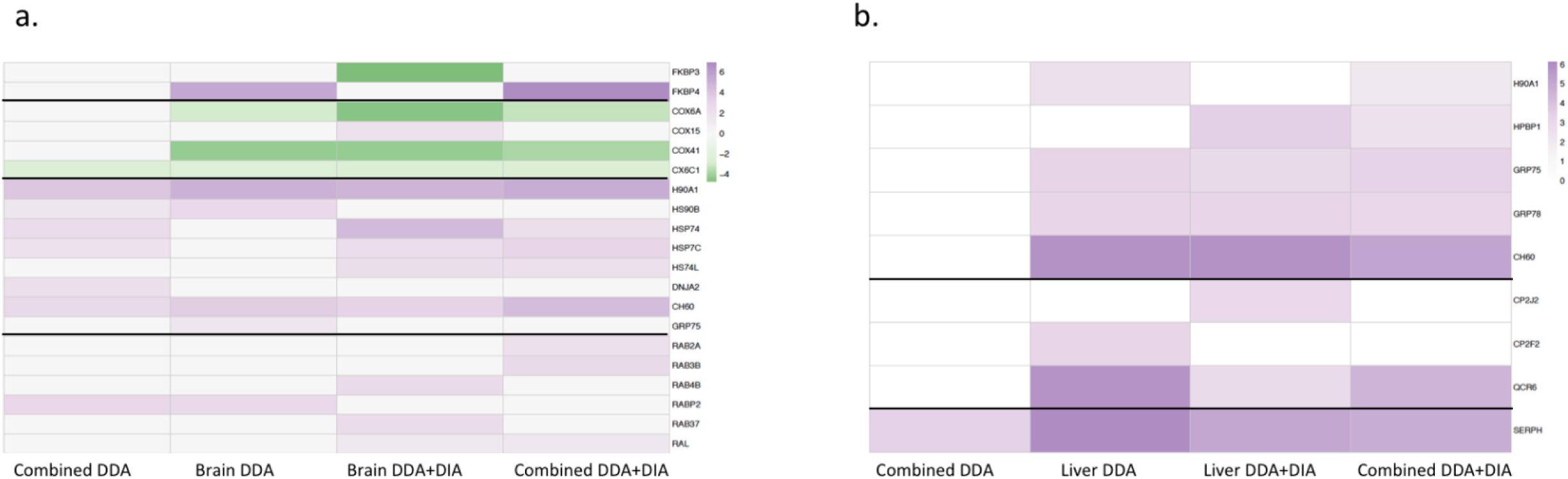
Heatmap of differentially expressed stress related proteins in different library comparisons, values refer to differences based on Tukey’s post hoc test after an ANOVA test of significance (FDR<0.05). a) those identified in brain DIA targeted analysis and b) those identified in liver analysis. Thick lines indicate separations between proteins related to shared functions.

## Discussion

Recent technological advances in proteomics have turned mass spectrometry into a mainstream analytical tool with ecological applications (Aebersold & Mann, 2016). SWATH/DIA-MS is one of the first to allow for a big data approach to proteomics via its ability to create large data matrices containing accurately measured proteins across various samples with minimal missing values (Aebersold & Mann, 2016). Here we are able to present one of the first applications of this method to examine ecologically relevant changes in protein expression in a non-model organism. As the limitation of this method is the quality of the spectral library we examined the utility of different library compositions to determine the most useful, both in overall protein identification and in the statistical analysis of differential expression (Krasny et al., 2018). We are able to show that while a tissue-specific library approach yields the highest data completion there is no loss of ecologically relevant information when using an organism level library, as long as that library contains study-specific data. This is especially important when working with small organisms, such as during developmental or larval stages where the biological material is usually limited. Further studies using non-model organisms can be confident that a whole organism approach to a spectral library will lead to successful differential expression analysis as well as promote future proteomic studies in the non-model species.

The spectral library generation is the most laboratory intensive and time-consuming part of the SWATH/DIA-MS methodology as it requires fractionation prior to mass spectrometry. However, once an organism-specific library is created future studies can use this reference cutting down on costs and lab time. Using a tissue-specific library led to a greater number of identifications at the peptide and protein level when querying samples of that same tissue, but when we compared the liver DIA samples to the brain-specific library or vice versa we discovered a significant loss in data completeness and protein identification. Therefore, unless one tissue will always be the focus in an organism, a comprehensive species-specific library will be more useful to future proteomic studies than a tissue-specific one.

We also undertook the first study to quantify the usefulness of adding study-specific DIA data to the spectral library in a non-model organism. A previous study suggested the utility of this method in lab curated cancer cells with positive results in increases of protein identifications (Gandhi et al., 2017). We discovered that the addition of study-specific DIA data in the creation of the spectral library, especially at the whole organism level, increased the identification of protein groups and peptides significantly. Creating a base organism level DDA spectral library then adding DIA specific runs to tailor that library to a specified experiment can lead to the best results of both a study-specific and whole organism library (Gandhi et al., 2017).

To validate the created spectral libraries, we used DIA samples from a complex experimental design with approximately twelve individuals from each of four different conditions. As the experiment was set up to identify the reaction of this fish to the dual climate stressors of ocean acidification and ocean warming, we searched the differentially expressed proteins for those related to common stress responses in fish. Proteins related to cellular stress were found in every brain and liver analysis using different library types. However, a more complex stress responses was identified using tissue-specific and study-specific (DIA+DDA) libraries. In particular we were only able to identify changes between the combined condition and the elevated temperature condition when using DDA+DIA libraries (Supplementary Data 1). This was a very small change consisting of less than five proteins but when using the DDA libraries the identification of any proteins between these conditions was lost. Proteins identified as part of the cellular stress response included up regulation of HSPs, a common indicator of stress related to abiotic and biotic factors in fish and other organisms (Basu et al., 2002; Huth & Place, 2016). We also identified several cytochrome proteins involved in oxidative and metabolic processes, and associated with thermal stress responses in salmon (Akbarzadeh et al., 2018). *SEPRH, FKBP*, and *RAS* related proteins have also been previously identified as significantly expressed in marine fish under stress (Chen et al., 2018; Evans & Somero, 2009; Liu et al., 2016). With these methods we are able to identify expected changes in relevant molecular pathways previously recognized as being associated with environmental stress responses in fish.

Our in-depth approach into the utility of different spectral libraries provides new insights into the use of SWATH/DIA-MS on non-model organisms and in complex experimental designs with high individual variation. We were able to identify up to 4,000 protein groups and up to 253 differentially expressed proteins using this method. Within the differentially expressed proteins we successfully identified many proteins related to stress responses in fish, confirming the ability of this method to identify ecologically relevant pathways when examining individuals exposed to varying environmental conditions. We provide the first proof-of-concept for the use of SWATH/DIA-MS in non-model organism from a wild, non-lab bred population. These results contribute to our better understanding of proteome changes in our own study organism, *A. polyacanthus*, as well as providing analytical data to encourage the use of this method in further ecological based proteome studies of any non-model organisms.

## Acknowledgments

We would like to thank Michael D. Jarrold, Megan J. Welch, and Philip L. Munday for their help with the live fish experiments and aid in collection of the fish tissues. We also thank the KAUST Proteomics Core Lab for their help throughout the sample analysis stage and the KAUST Integrative Systems Lab for all their support. This study was supported by the Office of Competitive Research Funds OSR-2015-CRG4-2541 from the King Abdullah University of Science and Technology (T.R., C.S., A.A.M.).

## Data accessibility

The mass spectrometry DDA and DIA proteomics data acquired during this experiment have been deposited to the ProteomeXchange Consortium via the PRIDE partner repository with the dataset identifier PXD017605 (Perez-Riverol et al., 2019).

## Author contributions

AAM, HZ, CS, and TR conceived of and designed the research. AAM and CS prepared the samples. AAM and HZ performed the experiments and processed the data. AAM analyzed the data. AAM wrote the manuscript with input and revisions from HZ, CS, and TR. All authors read and approved the manuscript.

