## Supplementary figures and images for "Probing SWATH-MS as a tool for proteome level quantification in a non-model fish"

### Supplementary Figure 1

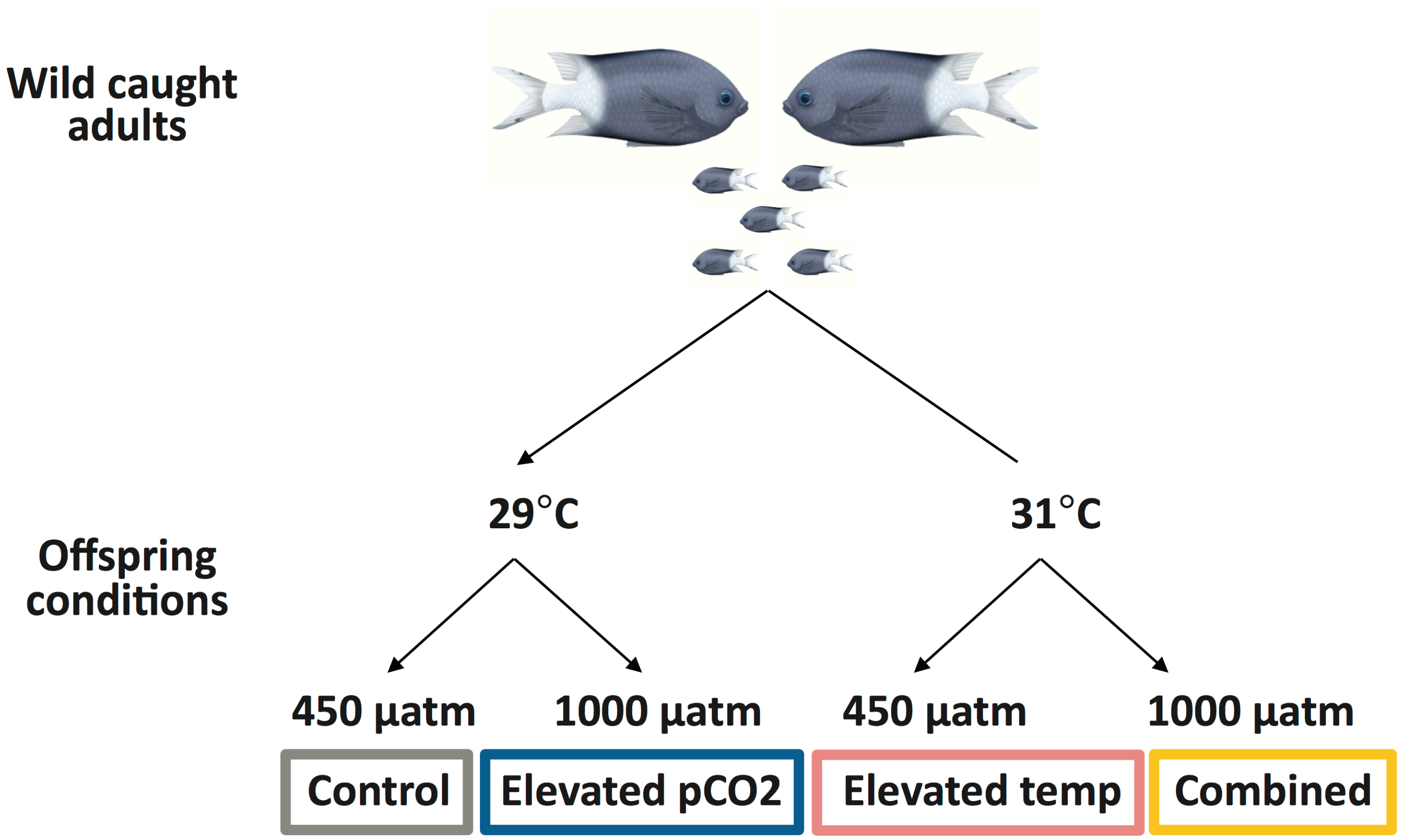

### Supplementary Figure 2

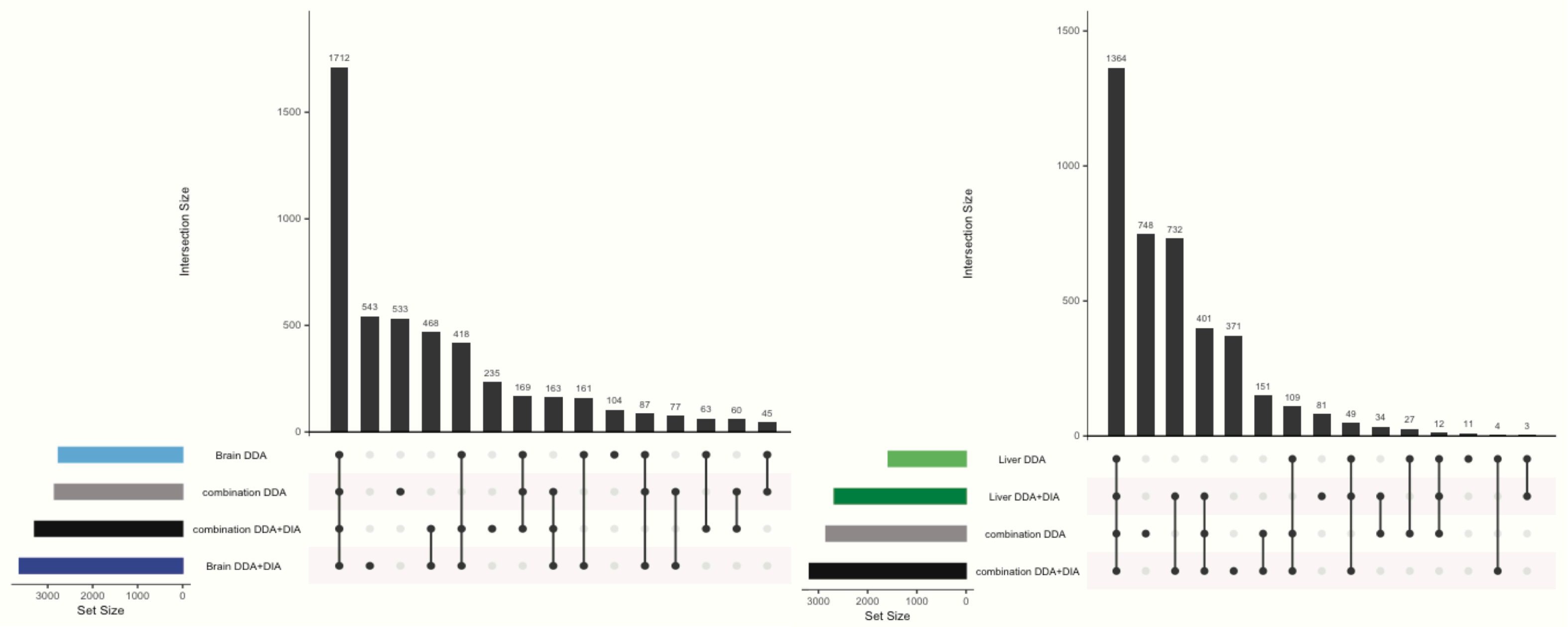
